# The neuronal GPCR OCTR-1 mediates temperature effects on longevity by regulating immune response genes in *C. elegans*

**DOI:** 10.1101/2021.05.07.443056

**Authors:** Shawndra Wibisono, Phillip Wibisono, Chia-Hui Chen, Jingru Sun, Yiyong Liu

**Author notes:** Correspondence (J.S.); (Y.L.).

## Abstract

Researchers have long known that many animals live longer in colder climates than in warmer climates. The inverse relationship between temperature and lifespan was traditionally explained using the rate of living theory, which suggests that higher temperatures increase chemical reaction rates, thus speeding up the aging process. However, recent studies have identified specific molecules, cells and signaling pathways involved in the longevity response to temperature, indicating that such a response is not simply thermodynamic but a regulated process. Here, we report that *Caenorhabditis elegans* lacking OCTR-1, a neuronal G protein-couple receptor for the neurotransmitter octopamine, had extended lifespan at warm temperature but shortened lifespan at cool temperature, indicating that OCTR-1 modulates the longevity response to both warm and cool temperatures. We further found that these responses are regulated by the OCTR-1-expressing, chemosensory ASH neurons. Transcriptomic analysis and functional assays revealed that OCTR-1 mediates temperature effects on longevity by regulating a subset of immune response genes. Our study provides cellular and molecular insights into the relationship between temperature and longevity, which could be useful for developing strategies to extend human lifespan in the midst of global warming.

## INTRODUCTION

It has been known for more than a century that many animals live longer in colder climates than in warmer climates (1). Aside from anecdotal evidence, the inverse effects of temperature on longevity were first recorded in poikilotherms including the fruit fly *Drosophila melanogaster* (2) and the nematode *Caenorhabditis elegans* (3), and later were also observed in homeotherms such as mice (4) and humans (5), although with some obscurity (e.g., females maintain higher body temperatures than males and yet females live longer (1)). Nonetheless, there is a clear trend for lower temperature being associated with longer lifespan and higher temperature with shorter lifespan across species. Most of these early studies interpretated the inverse effects of temperature on longevity using the rate of living theory or were in favor of this theory, which suggests that lower temperatures reduce chemical reaction rates, thus slowing down the aging process, whereas higher temperatures do the opposite (6). However, recent studies have identified specific molecules, cells and signaling pathways that are involved in the longevity response to temperature, indicating that such a response is not simply thermodynamic but a regulated process. Despite these progresses, the regulatory mechanisms underlying the temperature-longevity relationship remain unclear.

*C. elegans* has become an excellent model system for studying the relationship between longevity and temperature because of its short lifespan, invariant lineage, and genetic tractability (7). In the laboratory, the worm is routinely propagated under three temperatures (15, 20, and 25°C), and the inverse effects of these temperatures on lifespan have been well documented, i.e., a 75% increase in lifespan from a 5°C drop in temperature, consistent between 15-20 and 20-25°C (3,8). Using the *C. elegans* model system, researchers have revealed many molecular details about the temperature effects on longevity. For example, Lee and Kenyon found that the thermosensory AFD neurons and AIY interneurons are required for the worm’s lifespan at the warm temperature 25°C, and that such involvement is linked to the DAF-9/DAF-12 steroid-signaling pathway (9). The AFD neural circuit maintains longevity at warm temperatures through CRH-1, the *C. elegans* cyclic AMP-responsive element binding protein, and its transcriptional target FLP-6, both of which antagonize insulin signaling to affect longevity (10). At cold temperatures, TRPA-1, a cold-sensitive transient receptor potential (TRP) channel, detects temperature drop in the environment to extend lifespan through calcium signaling (11). Interestingly, in contrast to exposure to cold temperature during adulthood that prolongs *C. elegans* lifespan, low-temperature treatment during development reduces lifespan, and this differential temperature effect is also mediated by TRPA-1 (12). *C. elegans* lacking functional DAF-41, the worm’s homolog of p23 co-chaperone/prostaglandin E synthase-3, lives longer than wild-type worms at 25°C but shorter than wild-type at 15°C, indicating that DAF-41 modulates the longevity response to both cold and worm temperatures (8). Recently, we have demonstrated that in *C. elegans,* functional loss of NPR-8, a G protein-coupled receptor related to mammalian neuropeptide Y receptors, increases worm lifespan at 25°C but not at 20°C or 15°C, and that the lifespan extension at 25°C is controlled by the NPR-8-expressing AWB and AWC chemosensory neurons as well as AFD thermosensory neurons through regulating a subset of collagen genes (13). Our integrative transcriptomic analyses also revealed elevated metabolism at warm temperatures, providing a partial molecular basis for the rate of living theory (13). These findings confirmed that the temperature-induced longevity response is indeed a regulated process but also has a thermodynamic component. As the rate of living theory and the view of a regulated process are not mutually exclusive, warm temperatures may speed up the aging process in both thermodynamic and neurally regulated manners in living organisms. More studies are needed to dissect the molecular and cellular mechanisms underlying temperature-dependent longevity and the aging process.

Previously, we have shown that *C. elegans* lacking functional OCTR-1, a GPCR for the neurotransmitter octopamine (OA), exhibited an enhanced immune response and improved survival upon pathogen infection, and identified an OCTR-1-dependent neuroimmune regulatory circuit that includes the OCTR-1-expressing sensory neurons ASH and ASI, OA, and the OA-producing interneurons RIC (14–17). Under nonpathogenic conditions and the normal cultivation temperature 20°C, however, *octr-1* mutant worms had the same lifespan as wild-type worms (14,17). Here, we report that functional loss of OCTR-1 extended lifespan at 25°C but shortens it at 15°C, indicating that OCTR-1 modulates the longevity response to both warm and cool temperatures. We further found that these responses are regulated by the OCTR-1-expressing, chemosensory ASH neurons. Transcriptomic analysis and functional assays revealed that OCTR-1 mediates temperature effects on longevity by regulating a subset of immune response genes. Our study provides cellular and molecular insights into the relationship between temperature and longevity, which could be useful for developing strategies to extend human lifespan in the midst of global warming.

## RESULTS AND DISCUSSION

### OCTR-1 modulates the longevity response to both warm and cool temperatures

Previously, we have shown that compared to wild-type *C. elegans*, worms lacking functional OCTR-1 (*octr-1(ok371)* worms) had an enhanced immune response and improved survival upon infection with the human pathogen *Pseudomonas aeruginosa* strain PA14 (14–17). These phenotypes were also observed in other mutant strains that are deficient in the OCTR-1-dependent neuroimmune regulatory circuit, including *tbh-1(n3247)*, *ASH(-)[JN1713]*, and *RIC::TeTx* worms, indicating that the OCTR-1 circuit suppresses the innate immune response in *C. elegans* (14–17). These experiments were conducted at the normal propagation temperature of 20°C. At this temperature, however, *octr-1(ok371)* worms exhibited wild-type lifespan under nonpathogenic conditions (14,17). As *C. elegans* is also routinely propagated at 15°C and 25°C in the laboratory, we explored whether OCTR-1, or lack thereof, has any effects on lifespan under these temperatures. To this end, we compared the lifespan of *octr-1(ok371)* worms and wild-type worms at 15°C, 20°C or 25°C. Consistent with our previous studies (14,17), the two strains had similar lifespan at 20°C (Fig.1 A). Unexpectedly, *octr-1(ok371)* worms lived longer than wild-type worms at 25°C but had shorter-than-wild-type lifespan at 15°C (Fig.1 B and C). Because temperatures have inverse effects on longevity, i.e., lower temperatures are associated with longer lifespan and higher temperatures with shorter lifespan (1,3), our finding demonstrate that functional OCTR-1 is required for such temperature effects on longevity. In other words, OCTR-1 modulates the longevity response to both warm and cool temperatures.

**Fig. 1.**
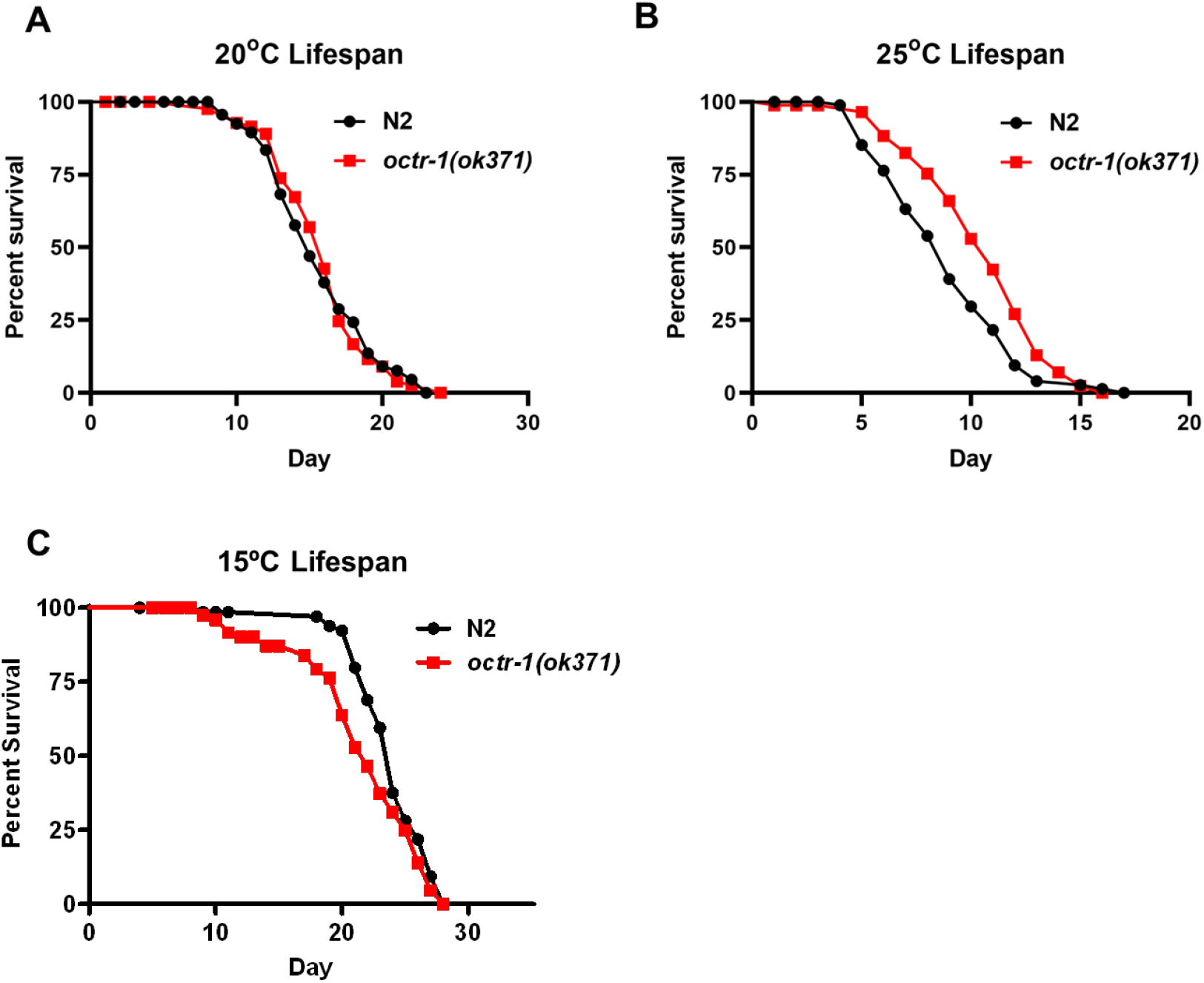
*octr-1(ok371)* worms had the same lifespan as wild-type worms at 20°C but lived longer at 25°C and shorter at 15°C. Wild-type N2 and *octr-1(ok371)* animals were grown on worm food *E. coli* OP50 and scored for survival over time at 20°C **(A)**, 25°C **(B)**, or 15°C **(C)**. The graphs are the representative results of three independent experiments. Each experiment included *n* = 90 adult worms per strain. *p* value represents the significance level of the mutants relative to the wild-type worms: *p* = 0.8410 for (A), *p* = 0.0018 for (B), and *p* = 0.0038 for (C).

### The OCTR-1-dependent longevity response to warm temperature is neurally regulated

We have previously demonstrated that OCTR-1 functions in the chemosensory neurons ASH to suppress the innate immune response in *C. elegans* by inhibiting the expression of immune genes (14). To determine whether ASH neurons are involved in the OCTR-1 neural circuit-regulated longevity response to warm temperature, we examined the lifespan of ASH-ablated worms *[ASH(-)* worms] and *ASH(-);octr-1(ok371)* double mutants along with wild-type and *octr-1(ok371)* worms at 25°C. As shown in Figure 2A, both *ASH(-)* worms and *ASH(-);octr-1(ok371)* double mutants lived significantly longer than wild-type N2 animals at 25°C, suggesting that the ASH chemosensory neurons play an important role in the OCTR-1-dependent regulation of longevity at warm temperature.

**Fig. 2.**
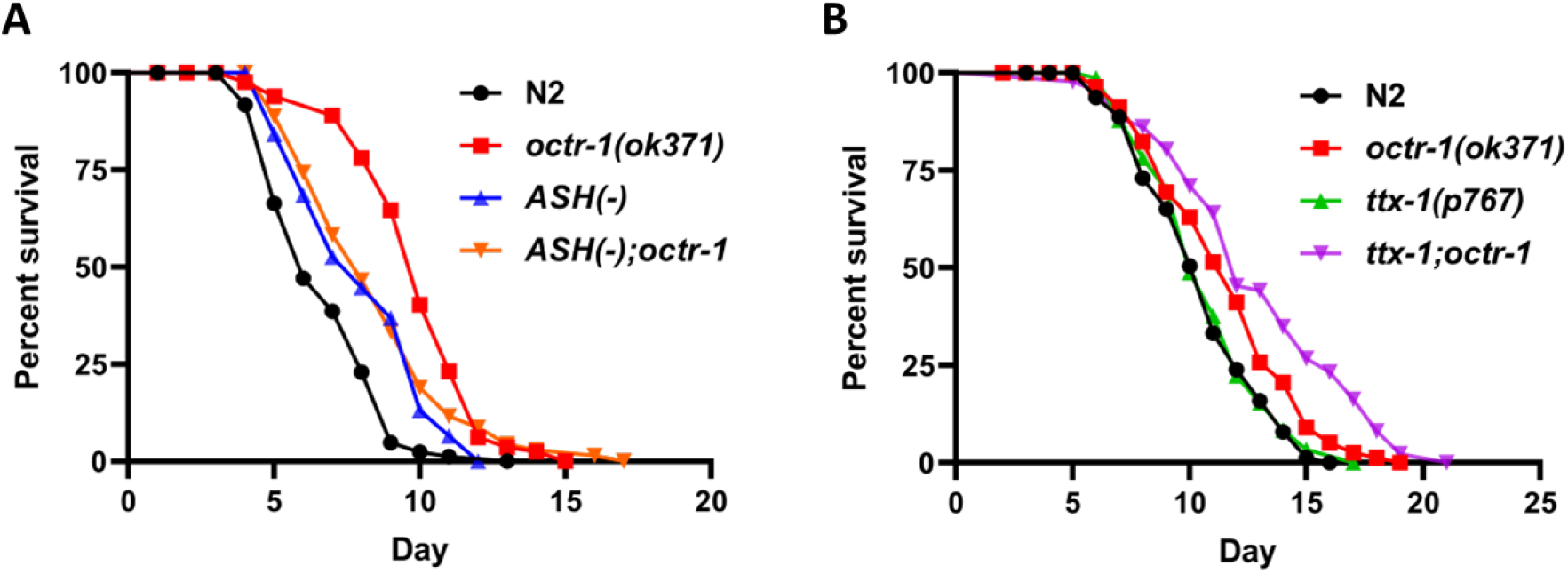
The ASH neurons, but not the AFD neurons, play an important role in the OCTR-1-dependent regulation of longevity at 25°C. (A) Wild-type N2, *octr-1(ok371), ASH(-)*, and *ASH(-);octr-1(ok371)* mutant animals were grown on *C. elegans* laboratory food *E. coli* OP50 and scored for lifespan over time at 25°C. The graph is a representative result of three independent experiments. Each experiment included n=90 adult animals per strain. *P* value represents the significance level of the mutants relative to the wild-type N2: *octr-1(ok371), p*<0.0001; ASH(-), *p*=0.0008; *ASH(-);octr-1(ok371), p*<0.0001. (B) Wild-type N2, *octr-1(ok371), ttx-1(p767)*, and *ttx-1(p767);octr-1(ok371)* mutant animals were grown on *C. elegans* laboratory food *E. coli* OP50 and scored for lifespan over time at 25°C. The graph is a representative result of three independent experiments. Each experiment included n=90 adult animals per strain. *P* value represents the significance level of the mutants relative to the wild-type N2: *octr-1(ok371), p*<0.0001; *ttx-1(p767), p*=0.1862; *ttx-1(p767);octr-1(ok371), p*<0.0001.

*C. elegans* senses temperature using the AFD thermosensory neurons. The thermosensory circuit for response to temperature changes has been well studied (18). To determine if the AFD thermosensory neurons are involved in the OCTR-1 neural circuit-regulated longevity at warm temperature, we examined the lifespans of AFD mutants *[ttx-1(p767)]* and *ttx-1(p767);octr-1(ok371)* mutants at 25°C. As shown in Figure 2B, the *ttx-1(p767);octr-1(ok371)* mutants lived significantly longer than wild-type N2 animals at 25°C, but no difference in lifespan was observed between AFD mutants and wild-type N2 animals. These results suggest that the AFD thermosensory neurons do not play an important role in the OCTR-1-dependent regulation of longevity at warm temperature. The role of the AFD neurons in sensing higher temperature at 25°C might be counterbalanced by an unidentified neuron(s) in the OCTR-1 neural circuit, which grants a future investigation.

### Cool temperature lowers the basal innate immune response at old age

To gain molecular insights into the OCTR-1-dependent temperature effects on longevity, we employed RNA sequencing to compare gene expression in young and old wild-type and *octr-1(ok371)* worms under the three growth temperatures. In this study, we define the young-adult stage as young age and the time when wild-type worms reached approximately 50% survival rate in lifespan assays as old age, which corresponds to Days 24, 14, and 9 of adulthood under the cultivation temperatures of 15°C, 20°C, and 25°C, respectively (Figs. 1 and S1). Five replicates of worms under each temperature condition were collected at young and old ages for RNA-seq analysis. Detailed sampling information is listed in Table 1. Total RNA was extracted from the collected worms, and RNA samples were submitted to the Washington State University Genomics Core for RNA-seq analysis.

**Table 1.** Grouping of RNA-seq samples.

| Sample Group | Strain | Propagation Temperature | Age* |
| --- | --- | --- | --- |
| 1 | *N2* | 15°C | young |
| 2 | *N2* | 15°C | old |
| 3 | *N2* | 20°C | young |
| 4 | *N2* | 20°C | old |
| 5 | *N2* | 25°C | young |
| 6 | *N2* | 25°C | old |
| 7 | *octr-1(ok371)* | 15°C | young |
| 8 | *octr-1(ok371)* | 15°C | old |
| 9 | *octr-1(ok371)* | 20°C | young |
| 10 | *octr-1(ok371)* | 20°C | old |
| 11 | *octr-1(ok371)* | 25°C | young |
| 12 | *octr-1(ok371)* | 25°C | old |

\* In this study, young age was defined as the young-adult stage, and old age was defined as the time when wild-type worms reached approximately 50% survival rate in lifespan assays, which corresponds to Days 24, 14, and 9 of adulthood under the cultivation temperature 15°C, 20°C, and 25°C, respectively. *octr-1(ok371)* worms were collected at the same biological ages as wild-type worms at these temperatures.

To examine how cool temperature affects gene expression, we compared the expression profiles of old wild-type worms grown at 15°C with those grown at 20°C. To this end, 14-day-old adults grown at 15°C and 9-day-old adults grown at 20°C were compared (sample group 2 vs. 4 in Table 1), as we considered these worms biological age-matched because at their respective ages, both worms had a 50% survival rate (Fig. 1). Our RNA-seq results showed that 1,010 genes were upregulated while 1,182 genes were downregulated at least 2-fold in worms grown at 15°C relative to those grown at 20°C. Gene ontology (GO) analysis of the upregulated genes identified enriched GO terms involving intraciliary transport and non-motile cilium assembly, the biological significance of which is not clear (Fig. 3A). GO analysis of the downregulated genes revealed eight classes of GO terms with “innate immune response” being the most significantly enriched (Fig. 3). The enriched innate immune response resulted from the downregulation of 95 immune genes encoding markers of immunity, such as CUB-like domain-containing proteins, ShK toxin domain-containing proteins, C-type lectin, lysozymes, glutathione s-transferease, saposin-like proteins, and others (Table S1). Our data suggest that cool temperature lowers the basal innate immune response in *C. elegans.* Interestingly, independent studies showed that knocking down some of these immune genes by RNAi, such as *dod-24*, *dod-22*, *clec-186*, *K04A8.1* (*dre-1*), *dod-17*, and *dod-19*, extended worm lifespan at 20°C (19–21), indicating that a low basal immune response promotes longevity.

**Fig. 3.**
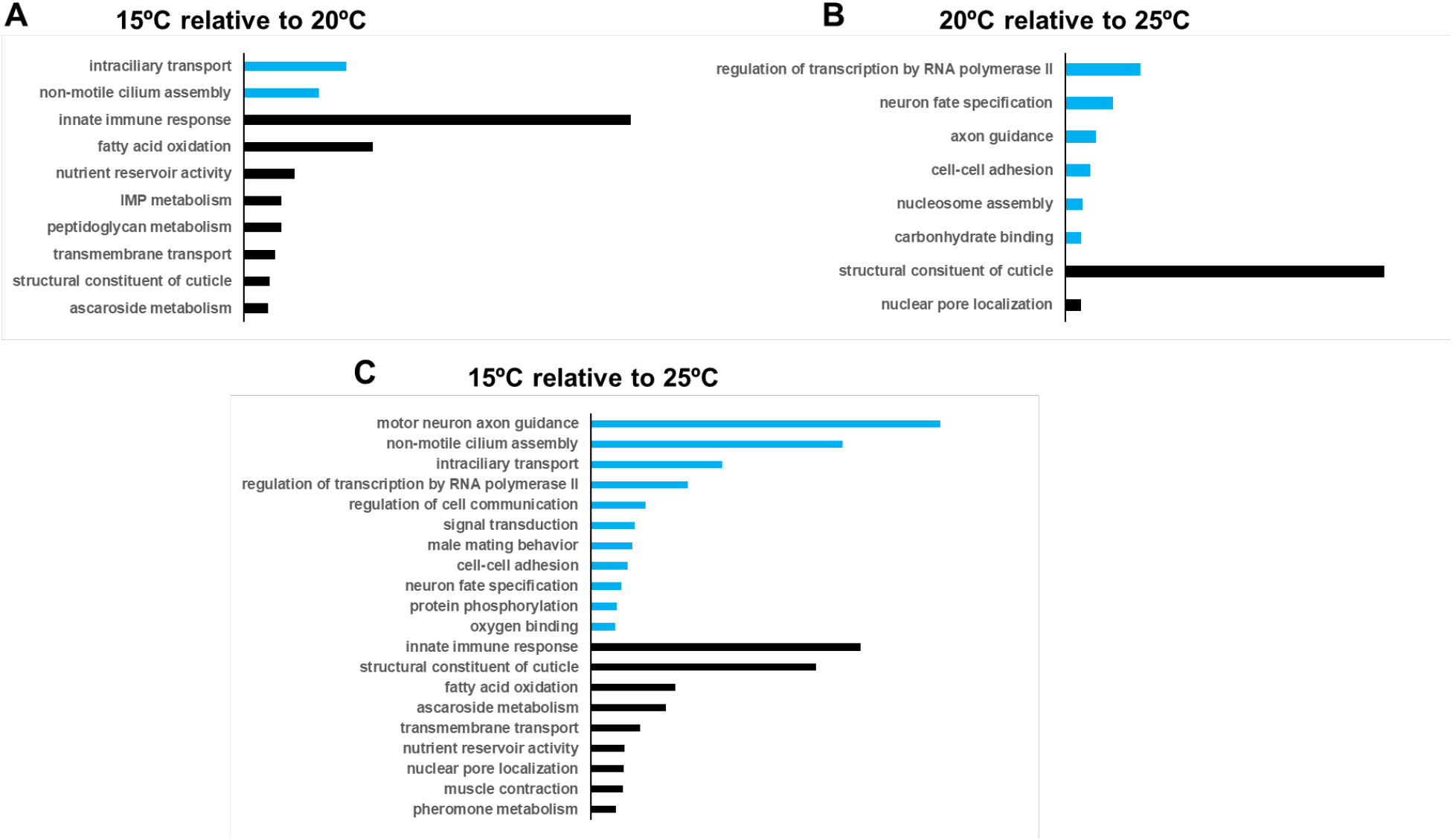
GO analyses revealed that cool temperature lowers the basal innate immune response at old age. GO analyses were performed with regulated genes in old wild-type worms grown at 15°C relative to those grown at 20°C **(A)**, grown at 20°C relative to 25°C **(B)**, or grown at 15°C relative to 25°C **(C)**. GO terms were categorized into different classes based on the PANTHER Classification System and presented in bar graphs. The length of each bar represents the significance of the enrichment based on FDR values. Blue bars, enriched GO term classes from upregulated genes; black bars, enriched GO term classes from downregulated genes.

The above-described analysis linked a decreased basal immune response to the lower temperature between 15°C and 20°C. We then asked whether this is also the case in the comparisons between 20°C and 25°C (sample group 6 vs. 4 in Table 1) as well as between 15°C and 25°C (sample group 2 vs. 6 in Table 1). GO analysis of the regulated genes in the latter two comparisons revealed that an enriched immune response was associated with 15°C between 15°C and 25°C but not with 20°C in the comparison between 20°C and 25°C (Fig. 3), demonstrating that a low basal immune response is specifically associated with 15°C but not necessarily associated with any lower temperature between two different temperatures. Taken together, our data suggest that a low basal innate immune response is cool temperature-specific, which likely contribute to the extended lifespan under cool temperatures.

### Functional loss of OCTR-1 alters signal transduction and the expression of immune and collagen genes

Previously, we have shown that *C. elegans* lacking OCTR-1 exhibited improved survival against pathogen infection due to the increased expression of immune genes and the concomitant enhancement of immunity (14–17). To determine how OCTR-1, or the lack thereof, impacts gene expression in *C. elegans* under nonpathogenic conditions, we compared the expression profiles of *octr-1(ok371)* worms grown at 20°C with those of wild-type controls at young or old age. At the young-adult stage, 790 genes were upregulated and 60 genes were downregulated at least 2-fold in the mutants relative to wild-type worms (sample group 9 vs. 3 in Table 1). While GO analysis of the downregulated genes did not yield any significantly enriched GO terms, a similar analysis of the upregulated genes identified three classes of GO terms, including protein (de)phosphorylation, structural constituent of cuticle, and innate immune response (Fig. 4A). These data indicates that functional loss of OCTR-1 alters signal transduction through protein phosphorylation and/or dephosphorylation as well as increases the basal innate immune response, the latter of which is consistent with our previous observation that inactivation of the OCTR-1 neuroimmune regulatory circuit causes elevated basal expression of immune genes under nonpathogenic conditions (17). Interestingly, the *octr-1* mutation also led to the enrichment of structural constituent of cuticle, which resulted from the upregulation of 32 cuticular collagen genes (Fig. 4A and Table S2). Cuticular collagens have been implicated in lengthening *C. elegans* lifespan under stress such as warm temperatures, pathogen infection, and oxidative stress (13,22,23). Upregulation of these collagen genes may also contribute to *octr-1(ok371)* worms’ enhanced resistance to pathogen infection and warm temperature.

**Fig. 4.**
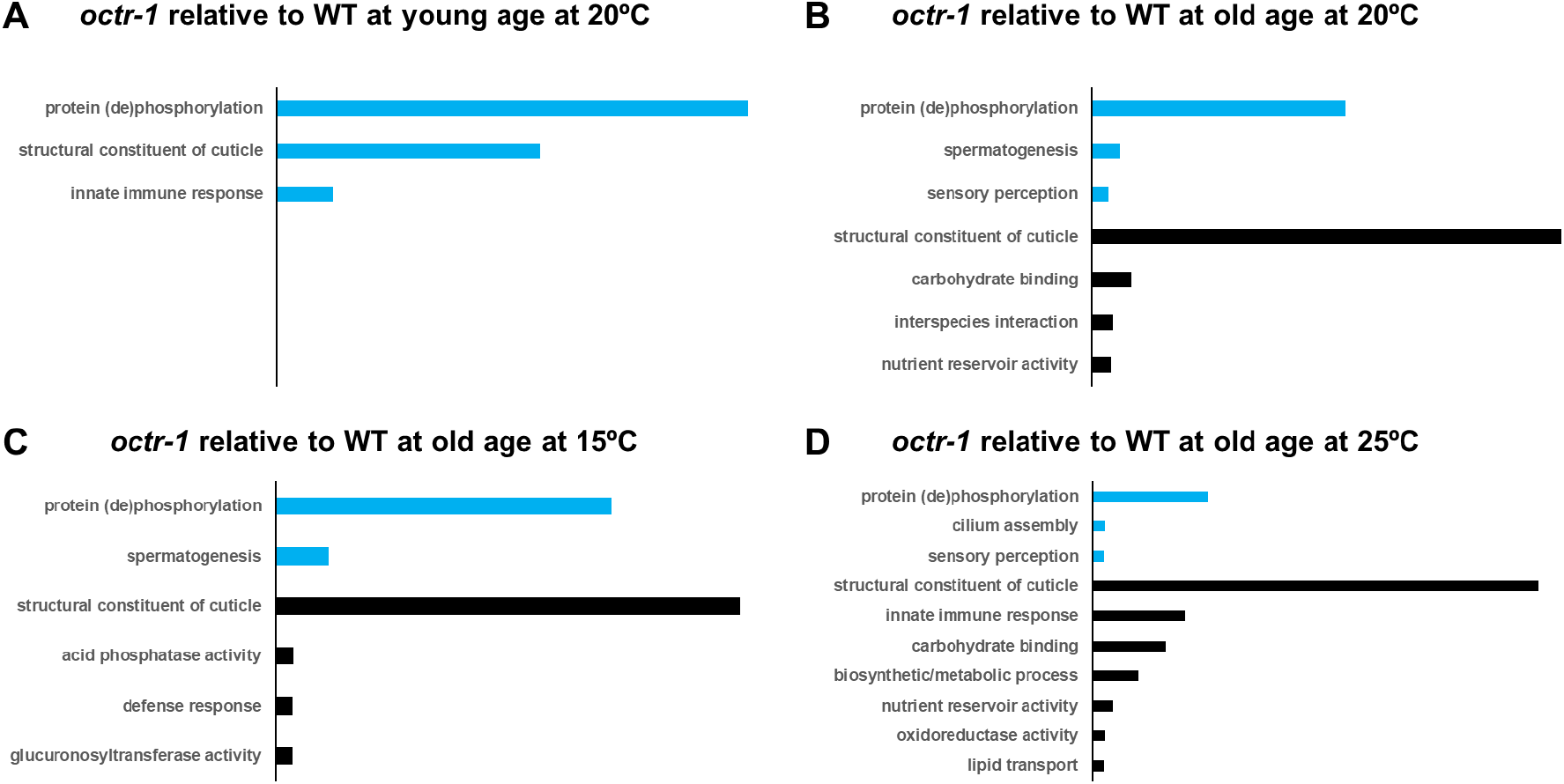
GO analyses revealed that OCTR-1 regulates protein (de)phosphorylation, collagen genes, and immune genes. GO analyses were performed with regulated genes in *octr-1(ok371)* worms relative to wild-type worms at young age at 20°C **(A)**, at old age at 20°C **(B)**, at old age at 15°C **(C)**, or at old age at 25°C **(D)**. GO terms were categorized into different classes based on the PANTHER Classification System and presented in bar graphs. The length of each bar represents the significance of the enrichment based on FDR values. Blue bars, enriched GO term classes from upregulated genes; black bars, enriched GO term classes from downregulated genes.

At the old age, GO analysis of the upregulated genes in *octr-1(ok371)* worms relative to wild-type worms (sample group 10 vs. 4 in Table 1) identified three classes of GO terms, with protein (de)phosphorylation being the most significantly enriched (Fig. 4B). This result indicates that OCTR impacts signal transduction through protein (de)phosphorylation at old age, like the above-described OCTR-1’s function at the young-adult stage, suggesting that such a function is independent of aging. GO analysis of the downregulated genes identified four classes of GO terms, with structural constituent of cuticle being the most significantly enriched (Fig. 4B). Downregulation of 45 cuticular collagen genes contributes to such an enrichment in the mutants (Table S3), indicating that at old age, OCTR-1 upregulates the expression of collagen genes in wild-type worms. This result is surprising and is in direct contrast with OCTR-1’s function at young age when it suppresses collagen genes (Fig. 4A). More strikingly, 16 of the 32 collagen genes that changed expression at young age were also regulated at old age (compare Table S2 with Table S3). These results suggest that the *octr-1* mutation could have beneficial effects on fitness in early life but are detrimental in later life, as suggested by George Williams’ antagonistic pleiotropic theory (24).

### OCTR-1 mediates the longevity response to warm temperature by regulating immune genes

Like many other species, *C. elegans* shows deceased lifespan at warm temperatures ((3) also Fig. 1). In our study, we observed that worms lacking functional OCTR-1 extended the short lifespan at 25°C (Fig. 1B), indicating that the *C. elegans* longevity response to warm temperature requires OCTR-1. To investigate the molecular basis underlying this phenomenon, we compared the gene expression profiles of old *octr-1(ok371)* worms to wild-type control worms grown at 25°C and found that 2,567 genes were upregulated while 1,168 genes were downregulated at least 2-fold in the mutants relative to the controls. GO analysis of the upregulated genes identified three classes of GO terms, including protein (de)phosphorylation, cilium assembly, and sensory perception, indicating alterations in signal transduction in the mutant worms (Fig. 4D). A similar analysis of the downregulated genes revealed the enrichment of seven classes of GO terms, which include innate immune response that was not enriched at the normal growth temperature 20°C (Fig. 4D). Fifty-one downregulated immune genes contributed to the enrichment of the immune defense response at 25°C (Table S4). These results suggest that *octr-1(ok371)* worms have a lower basal immune response than wild-type worms at old age at 25°C, which could explain the mutants’ extended lifespan at the warm temperature. These data also indicate that OCTR-1 increases the expression of immune genes in wild-type worms at old age at 25°C, which is opposite to its function at young age at 20°C, where it suppresses immune gene expression (Fig. 4A), suggesting that how OCTR-1 regulate immune genes is context-dependent.

To investigate whether the OCTR-1-controlled immune genes play any roles in lifespan extension at 25°C, we individually inactivated three top regulated genes listed in Table S4 (*acdh-1*, lys-5, and *lys-4*) by RNAi in wild-type worms and then assayed their lifespan at 25°C. As demonstrated in Fig. 5, RNAi of each of these genes significantly extended the lifespan of wild-type worms at 25°C, recapitulating the lifespan phenotype of *octr-1(ok371)* worms at the warm temperature. These results indicate that OCTR-1-controlled immune genes play a critical role in lifespan regulation at the warm temperature, and that decreased expression of immune genes in *octr-1(ok371)* worms reflected a low basal immune response in the mutants that likely led to their extended lifespan. This notion is consistent with the above-described observation with cool temperature in which a low basal immune response extends lifespan. Taken together, our data suggest that OCTR-1 mediates the longevity response to warm temperature by regulating immune genes.

**Fig. 5.**
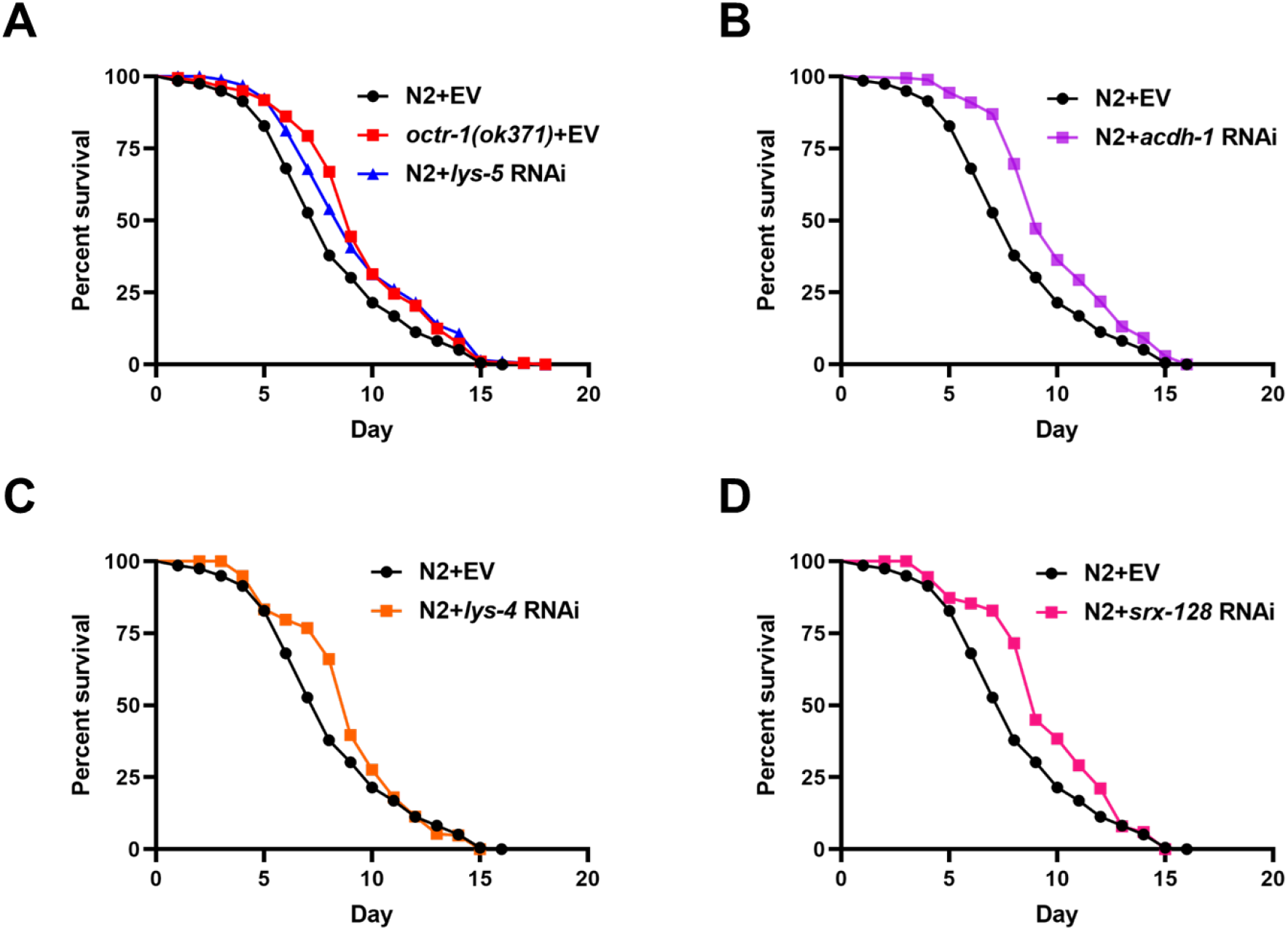
The OCTR-1-regulated immune defense genes contribute to the improved lifespan of *octr-1(ok371)* animals at 25°C. Wild-type N2 and *octr-1(ok371)* animals grown on dsRNA for several top-downregulated genes or empty vector (EV) control were exposed to *C. elegans* laboratory food *E. coli* OP50 and scored for lifespan over time at 25°C. The graphs are the combined results of three independent experiments. Each experiment included *n*=60 adult animals per strain. *P* value represents the significance level of the mutants relative to the wildtype N2+EV: (A) *octr-1(ok371)*+EV, *p*<0.0001; N2+*lys-5* RNAi, *p*=0.0007; (B) N2+*acdh-1* RNAi, *p*<0.0001; (C) N2+*lys-4* RNAi, *p*=0.0296; (D) N2+*srx-128* RNAi, *p*=0.0001.

## MATERIALS AND METHODS

### Nematode strains

The following *C. elegans* strains were cultured under standard conditions and fed *Escherichia coli* OP50. Wild-type N2 were *C. elegans* Bristol N2. The *octr-1(ok371), ttx-1(p767)*, and ASH(-) [JN1713] strains were obtained from the *Caenorhabditis elegans* Genetics Center (University of Minnesota, Minneapolis, MN). The *ASH(-);octr-1(ok371)* and *ttx-1(p767);octr-1(ok371)* mutant strains were constructed using standard genetic techniques.

### Bacterial strain

*E. coli* strain OP50 were grown in Luria-Bertani (LB) broth at 37°C.

### Lifespan assay

*C. elegans* wild-type *N2* animals and mutant animals were maintained as hermaphrodites at 20°C, grown on modified nematode growth medium (NGM) agar plates (0.35% instead of 0.25% peptone). Bacterial lawns used for the lifespan assays were prepared by placing a 50μL drop of an overnight fresh culture of *E. coli* OP50 on each 3.5cm plate. Plates were incubated at 37°C for 16 hours, cooled down to room temperature, and then seeded with synchronized 65-hour-old young adult animals. These worms were prepared by undergoing two rounds of egg-laying synchronization before they were exposed to *E. coli* OP50 for lifespan assays. The lifespan assays were performed at 20°C or 25°C. Live animals were transferred daily to fresh plates. Animals were scored at the times indicated and were considered dead when they failed to respond to touch.

### RNA sequencing

Five replicates of four groups of RNA samples [*octr-1(ok371)*] and wild-type N2 animals grown on *E. coli* OP50 for 14 days at 20°C or 9 days at 25°C were collected and submitted to the Genomics Core for RNA-seq analysis. The integrity of total RNA was assessed using Fragment Analyzer (Advanced Analytical Technologies, Ankeny, IA) with the High Sensitivity RNA Analysis Kit. RNA quality numbers (RQNs) from 1 to 10 was assigned to each sample to indicate its integrity or quality. “10” stands for a perfect RNA sample without any degradation, whereas “1” marks a completely degraded sample. RNA samples with RQNs ranging from 8 to 10 were used for RNA library preparation with the TruSeq Stranded mRNA Library Prep Kit (Illumina, San Diego, CA). Briefly, mRNA was isolated from 2.5 μg of total RNA using poly-T oligo attached to magnetic beads and then subjected to fragmentation, followed by cDNA synthesis, dA-tailing, adaptor ligation and PCR enrichment. The sizes of the RNA libraries were assessed by Fragment Analyzer with the High Sensitivity NGS Fragment Analysis Kit. The concentrations of the RNA libraries were measured using a StepOnePlus Real-Time PCR System (ThermoFisher Scientific, San Jose, CA) with the KAPA Library Quantification Kit (Kapabiosystems, Wilmington, MA). The libraries were diluted to 2 nM with RSB (10 mM Tris-HCl, pH8.5) and denatured with 0.1 N NaOH. Eighteen pM libraries were clustered in a high-output flow cell using HiSeq Cluster Kit v4 on a cBot (Illumina). After cluster generation, the flow cell was loaded onto a HiSeq 2500 for sequencing using a HiSeq SBS kit v4 (Illumina). DNA was sequenced from both ends (paired-end) with a read length of 100 bp. The raw BCL files were converted to FASTQ files using the software program bcl2fastq2.17.1.14. Adaptors were trimmed from the FASTQ files during the conversion. On average, 40 million reads with a read length of 2x 100 bp were generated for each sample. RNA-seq data (FASTQ files) were aligned to the *C. elegans* reference genome (ce10, UCSC) using HISAT2. Gene expression quantification and differential expression were analyzed using HTSeq and DESeq2.

### RNA interference

RNAi was conducted by feeding L2 or L3 larval *C. elegans E. coli* strain HT115(DE3) expressing double-stranded RNA (dsRNA) that was homologous to a target gene (25,26). *E. coli* with the appropriate dsRNA vector were grown in LB broth containing ampicillin (100μg/mL) at 37°C for 16 hours and plated on modified NGM plates containing 100μg/mL ampicillin and 3mM isopropyl β-D-thiogalactoside (IPTG). The RNAi-expressing bacteria were allowed to grow for 16 hours at 37°C. The plates were cooled down before the L2 or L3 larval animals were placed on the bacteria. The animals were incubated at 20°C for 24 hours or until the animals were 65 hours old. *unc-22* RNAi was included as a positive control in all experiments to account for RNAi efficiency. Clone identity was confirmed by sequencing at Eton Bioscience Inc. (San Diego, CA).

### Statistical analysis

Lifespan curves were plotted using GraphPad PRISM (version 9) computer software. Lifespan was considered different from the appropriate control indicated in the main text when *P* <0.05. PRISM uses the product limit or Kaplan-Meier method to calculate survival fractions and the log-rank test, which is equivalent to the Mantel-Haenszel test, to compare survival curves. All experiments were repeated at least three times, unless otherwise indicated.

## ACKNOWLEDGEMENTS

We would like to thank the *Caenorhabditis* Genetics Center (CGC) for providing several worm strains used in this study. CGC is funded by the NIH Office of Research Infrastructure Programs (P40 OD010440).

## AUTHOR CONTRIBUTIONS

S.W., P.W., C.C., J.S., and Y.L. designed and performed experiments and analyzed data. J.S. and Y.L. wrote the paper.

## DATA AND MATERIALS AVAILABILITY

The RNA-seq data will be deposited in NCBI’s SRA database through the GEO. The processed gene quantification files and differential expression files will be deposited in the GEO. All data needed to evaluate the conclusions in the paper are present in the paper. Additional data related to this paper may be requested from the authors. The *C. elegans* strains and plasmids constructed by us and primer sequences used for their construction are available upon request.

